# Deep Learning for Predicting 16S rRNA Gene Copy Number

**DOI:** 10.1101/2022.11.26.518038

**Authors:** Jiazheng Miao, Tianlai Chen, Mustafa Misir, Yajuan Lin

## Abstract

**Background:** Culture-independent 16S rRNA gene metabarcoding is a commonly used method in microbiome profiling. However, this approach can only reflect the proportion of sequencing reads, rather than the actual cell fraction. To achieve more quantitative cell fraction estimates, we need to resolve the 16S gene copy numbers (GCN) for different community members. Currently, there are several bioinformatic tools available to estimate 16S GCN, either based on taxonomy assignment or phylogeny.

**Method:** Here we develop a novel algorithm, Stacked Ensemble Model (SEM), that estimates 16S GCN directly from the 16S rRNA gene sequence strings, without resolving taxonomy or phylogeny. For accessibility, we developed a public, end-to-end, web-based tool based on the SEM model, named Artificial Neural Network Approximator for 16S rRNA Gene Copy Number (ANNA16).

**Results:** Based on 27,579 16S rRNA gene sequence data (rrnDB database), we show that ANNA16 outperforms the most commonly used 16S GCN prediction algorithms. The prediction error range in the 5-fold cross validation of SEM is completely lower than all other algorithms for the 16S full-length sequence and partially lower at 16S subregions. The final test and a mock community test indicate ANNA16 is more accurate than all currently available tools (i.e., rrnDB, CopyRighter, PICRUSt2, & PAPRICA). SHAP value analysis indicates ANNA16 mainly learns information from rare insertions.

**Conclusion:** ANNA16 represents a deep learning based 16S GCN prediction tool. Compared to the traditional GCN prediction tools, ANNA16 has a simple structure, faster inference speed without precomputing, and higher accuracy. With increased 16S GCN data in the database, future studies could improve the prediction errors for rare, high-GCN taxa due to current under sampling.

## 1 INTRODUCTION

Culture-independent 16S rRNA metabarcoding is a molecular biology technique that applies DNA sequencing technology to the prokaryotic 16S rRNA gene to identify and quantify the microbial communities present in a sample [1]. This technique has been established as the gold standard for inferring organismal phylogeny and microbial ecology for over 20 years [2]. However, a major limitation of 16S rRNA metabarcoding is that next-generation sequencing estimates the proportion of 16S rRNA gene read counts, rather than the true cell counts. Since it is typical for prokaryotes to have more than one copy of the 16S rRNA gene per cell (1 - 21 copies/genome) [3], the proportion of 16S rRNA gene read counts (also known as relative 16S abundance) does not accurately reflect the true composition of a microbial community [4, 5]. To address this issue and accurately represent the profile of the microbiota, the proportion of 16S rRNA gene read counts needs to be weighted by the inverse of 16S gene copy number (GCN). This step yields a corrected microbial composition that represents the proportion of cell counts of each lineage in the community [4].

To estimate 16S GCN, researchers typically rely on experimental methods such as whole genome sequencing [3] or competitive PCR [6]. However, these methods can be expensive and/or culture-dependent, and thus experimental GCN estimates are only available for a limited number of species. To overcome this limitation, several bioinformatics tools have been recently developed to predict the GCN of unmeasured species from measured species. Although there are exceptions [7], the basic assumption is that 16S GCN is correlated with phylogenetic distance across species [4, 8], i.e., 16S GCN variation is relatively small among closely related species and increases with phylogenetic distance. This allows 16S GCN to be estimated either from taxonomy or phylogeny [4, 8, 9]. For instance, rrnDB [3] is a well-cited tool that corrects 16S GCN based on taxonomy. It calculates the mean 16S GCN of a taxon from the means of its sub-taxa. Another example is PICRUSt2 [10], one correction tool that constructs a phylogenetic tree of the 16S sequence and estimates the GCN of the unmeasured species from its close measured relatives.

Deep learning, as a popular machine learning strategy, has been recently applied to learn complex patterns from biological sequencing data [11, 12]. In this study, we explore deep learning as a novel approach for predicting 16S GCN. Deep learning is a branch of machine learning which uses many layers of Artificial Neural Networks (ANN) to learn complex representations of data [13]. The concept of ANN is inspired by the way a human brain learns and processes information through synaptic interactions between neurons [14]. An ANN is a computational model that is constituted by multiple interconnected processing elements— artificial neurons [14]. Neurons are utilized in groups as the layers of a network. Each artificial neuron receives and processes input with a non-linear function, known as the activation function [15]. Collectively, one ANN model learns to solve a specific task by capturing non-linear trends and relationships from the presented data. Beyond these capabilities, ANNs are even competent in disclosing relationships or mechanisms that are still unknown to scientists [16]. Deep learning gets its name by having more than one (hidden) layer connecting the input and output units. Having deeper networks refer to representing complex input-output relationships while accommodating a large amount of data. In that sense, high-dimensional biological data is a popular target for deep learning. For example, various deep learning architectures, like the Convolutional Neural Network, and Recurrent Neural Network, have been applied to microbiome research, including metagenomic-based taxonomy classification [17], 16S rRNA classification [18], host phenotype prediction [19], disease prediction [20], and microbial community predication [21]. Furthermore, deep learning algorithms have been developed to conduct data mining for antibiotic resistance genes [22], antimicrobial peptides [23], and microbial source [24].

With the great potential of extracting biological information from DNA sequences demonstrated by the previous studies, this study explores the possibility of estimating GCN directly from the 16S DNA sequence through deep learning. To the best of our knowledge, no deep learning-based method has been utilized for 16S GCN prediction yet. A Stacked Ensemble Model (SEM) is first trained on the 16S full-length sequences and compares the performance to the taxonomy-based and phylogeny-based algorithms. The sequences and GCN data are retrieved from the ribosomal RNA operon copy number database (rrnDB), a publicly available, curated resource for copy number information for bacteria and archaea [3]. Because microbial studies tend to target one to three adjacent hypervariable regions in 16S rRNA [25], this study also trains deep learning models on different combinations of the adjacent hypervariable regions in 16S rRNA and compares the performance with conventional algorithms. Finally, a deep learning-based bioinformatic tool for 16S GCN correction is developed. The tool is compared with existing correction tools to assess the performance. Shapley Additive exPlanations (SHAP), a game theory-based model explanation method [26], is applied to ANNA16 to reveal how this deep learning model understands the 16S rRNA gene sequences.

## 2 MATERIALS AND METHODS

### 2.1. Data Source and Preprocessing

16S rRNA gene sequences and the GCN data used to train deep learning models in this study were retrieved from rrnDB, version 5.7 [3]. The dataset includes a total of 20,277 16S gene sequences with unique accession numbers, containing both (+) strands and (-) strands.

Firstly, the orientation for all sequences in this dataset were identified using BLASTN algorithm (BLAST+, version 2.11.0) [27]. The first 50 bp of the *E. coli* 16S rRNA gene sequence (+) strand (accession number: GCF_002953035.1), a conserved region on 16S rRNA [28], were used as a template. Sequences with High-Scoring Segment Pairs (HSPs) that had sstart (subject start) > send (subject end) and evalue < 0.01 were identified as (+) strands, and those with sstart < send and evalue < 0.01 were identified as (-) strands. 389 sequences that did not meet these criteria were labeled as unrecognizable and excluded from the dataset. The reverse complement of these (-) strand sequences were then added to the (+) strand sub-dataset.

Next, in order to have uniform starting and ending positions, all 16S rRNA gene sequences in the database were further trimmed by HyperEx [29], using the universal full-length 16S primer 27F and primer 1492R (Table S1) [25, 30]. After that, 19,520 sequences remained in this full-length dataset, involving 1,331 genera, with an average sequence length of 1,498.72 nucleotides and a standard deviation of 21.41 nucleotides. 16S GCN ranges from 1 to 21 copies/genome (Mean = 5.26, SD = 2.74). In addition, seven commonly targeted subregions by metabarcoding, i.e., V1-V2, V1-V3, V3-V4, V4, V4-V5, V6-V8, and V7-V9 [25], were also extracted by HyperEx (primers listed in Table S1) to generate subregion datasets. The full-length and subregional datasets were used as the training datasets.

Additionally, during the course of this study, a new version rrnDB 5.8 were release on June 23rd, 2022, including 8,059 newly added sequences and their GCN. The added data involves 676 genera, and 219 of the genera that are not present in the version 5.7 dataset. The average sequence length is 1,499.79 nucleotides (SD = 21.67), and the range of 16S GCN is 1 - 20 copies/genome (Mean = 5.72, SD = 2.85). The newly published data were also processed by BLASTN and HyperEx and produced an independent final test dataset.

By conducting a five-fold cross-validation at each region, the initial GCN datasets were first split into training and test sets to compare the performance of SEM and other conventional algorithms. Data in the train datasets were used to pre-train a SEM model, and the model was tested at each region using the test datasets. Existing GCN tools were also applied to the test datasets and their performance were compared with that of SEM. Once an optimal GCN model was selected, the train datasets and test datasets were combined to train the final model. Finally, the independent final test dataset was used to evaluate the robustness and accuracy of the final model.

### 2.2. Method Comparison: Conventional GCN Algorithms and Existing Tools

There are two predominant CGN algorithms: taxonomy-based and phylogeny-based approaches.

Taxonomic aggregation (TA) is a commonly used taxonomy-based GCN algorithm [3, 31]. The algorithm calculates the mean 16S GCN of a taxon from the GCN means of its sub-taxa. To implement this algorithm, 16S sequences were first classified using the *assignTaxonomy* function in the DADA2 package in R (version 1.21.0) [32]. Two well-established 16S database, RDP (version 18) [33] and Greengenes (version 13.8) [34] were used as the reference for the *assignTaxonomy* function to determine whether the choice of classification database would impact GCN prediction performance. The algorithm was implemented in Python (version 3.8) with an object-oriented programming approach. The algorithm took the taxonomic and GCN data and treated every taxon as a Node object. Every Node is linked with its downstream Nodes (taxa), so eventually the Nodes formed a hierarchical tree where the *Prokaryote domain* is the root, and the species are the tips. Finally, the algorithm pre-computed the GCN of every Node by taking the mean of its sub-Nodes. When a query is given, the algorithm will search the hierarchical tree for the lowest taxon of the query lineage. If a query has unassigned taxon, it assigns the value of the query’s parent node to the query node as an estimation.

Phylogenetic algorithms were based on a pre-constructed tree of 16S rRNA gene sequences and the GCN of each ancestral node was estimated from its daughter species through weighing each labeled daughter node with phylogenetic distance to the parental node [35]. For method comparison, this study employed five phylogeny-based algorithms, including empirical probabilities (EP), subtree averaging (SA), phylogenetic independent contrasts (PIC) [36], Sankoff’s maximum-parsimony (MPR) [37], and weighted-squared-change parsimony (WSCP) [38]. To implement these algorithms, 16S rRNA gene sequences in the dataset were first aligned by MAFFT v7.490 [39] and then a 16S phylogenetic tree was constructed by FastTree 2.1.10 [40]. The five phylogeny-based GCN algorithms were then executed using the castor package in R [35], with modifications made from the R script in [9]. In particular, MPR was implemented by function *hsp_max_parsimony*, WSCP by function *hsp_squared_change_parsimony*, SA by function *hsp_subtree_averaging*, EP by function *hsp_empirical_probabilities*, and PIC by *hsp_independent_contrasts*, all configured to default parameters.

Apart from conventional GCN algorithms, the existing GCN tools include rrnDB [3], CopyRighter [4], PAPRICA [41], and PICRUSt2 [10] were also implemented for method comparison. rrnDB is a taxonomy-based tool using taxonomic aggregation, while the latter three are phylogeny-based tools. CopyRighter uses the PIC algorithm to pre-compute the copy number of all taxa in the Greengenes database [34] and makes predictions by searching the lineages of the queries in its pre-computed GCN table [4]. PAPRICA and PICRUSt2 first utilize EPA-ng [42] to place the queries on their built-in phylogenetic tree and then use subtree- averaging (PAPRICA) or maximum parsimony (PICRUSt2) to predict the GCN of the queries [10, 41].

Because rrnDB and CopyRighter output GCN corrected microbial community composition rather than 16S GCN values directly, this study extracted their precomputed tables (rrnDB version 5.7) to generate GCN values, searching the precomputed tables for assigned lineages. The taxonomy of full-length 16S sequences and all subregion sequences were generated by the *assignTaxonomy* function in the DADA2 package through RDP (version 18) and Greengenes (version 13.8, formatted by DADA2) [34] database. The assigned RDP lineages were searched in the rrnDB precomputed table, and Greengenes lineages were searched in CopyRighter precomputed table. As for PAPRICA and PICRUSt2, the DNA sequences in the test dataset were directly input to the tools. For PAPRICA, *paprica-run.sh* was called without additional parameters. For PTCRUSt2, *place_seqs.py* and *hsp.py* were called without additional parameters.

### 2.3. Development of ANNA16

#### 2.3.1. DNA sequence K-merization

The 16S rRNA gene sequences were first preprocessed with K-merization. An example of a set with three 6-mers is displayed in Figure 1a. The K-mers were converted into the number of counts in the set via the *CountVectorizer* function in the scikit-learn module (version 0.23.2) [43]. Subsequently, the K-mer counts and GCN data were used to train the SEM models.

**Figure 1.**
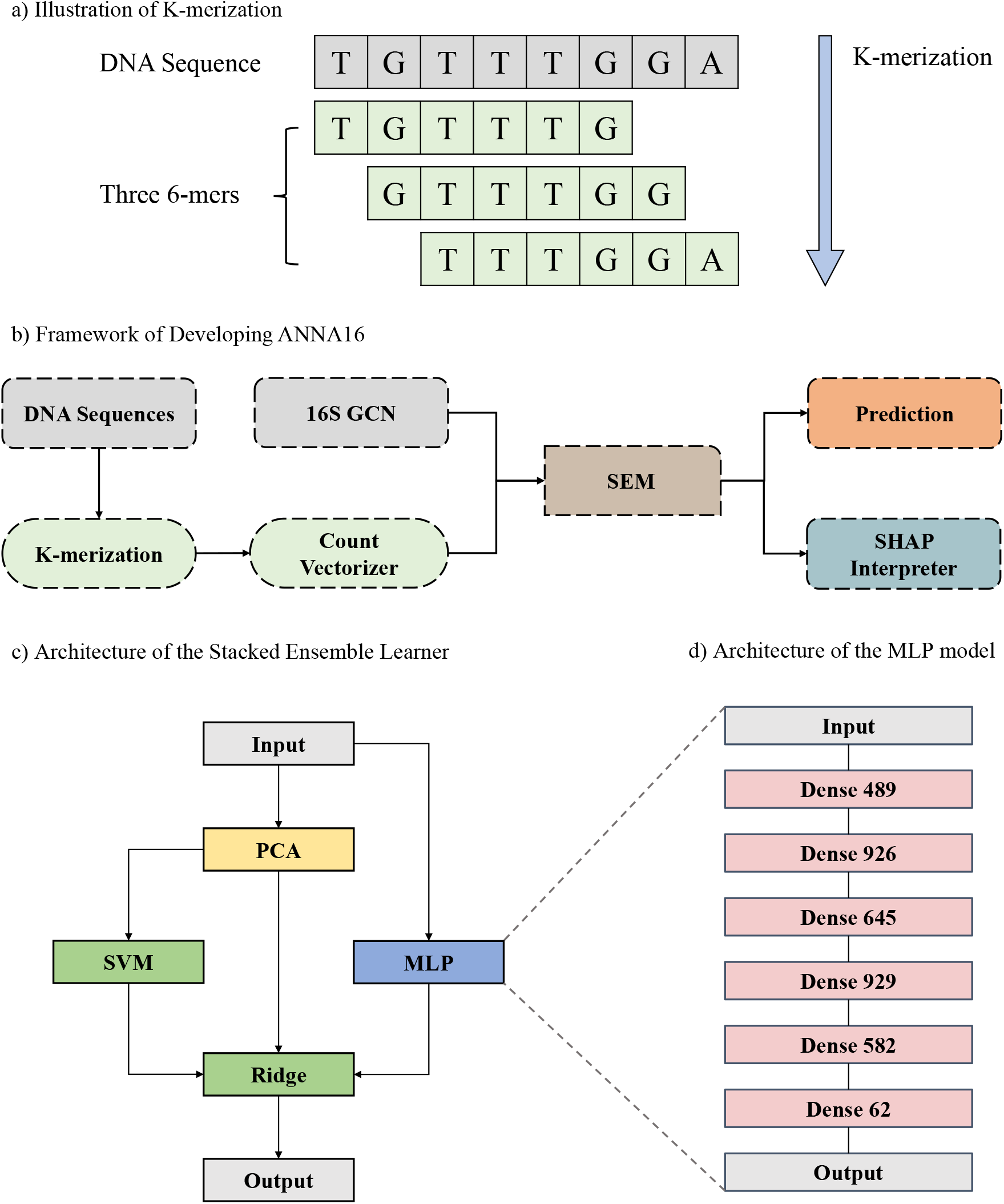
Illustration of the framework and architecture of the deep learning models. a) DNA sequence K-merization (K = 6). b) Framework for designing ANNA16. c) Architecture of the Stacked Ensemble Model (SEM). d) Architecture of the MLP model module inside the SEM.

After testing K from 3 to 7 on a prototype 3-layer Multi-layer Perceptron (MLP) model, the model performance was found to improve with increasing K. However, a larger K value required significantly more memory. Thus, the optimal K value was determined to be 6, striking a balance between good performance and manageable memory requirements.

#### 2.3.2. Model Construction

Based on 6-mers, the Stacked Ensemble Model (SEM) was constructed for 16S GCN prediction using the Tensorflow package (version 2.8.0) in Python (version 3.8). The initial dataset was split for training and testing by 3:1. The SEM model training was conducted on a Nvidia RTX 3090 GPU. This SEM method involves training multiple base models on the available data, after which a meta-model is used to aggregate the outputs of all base-models and make the final prediction. Typically, the stacked model performs better than any single base model [44, 45].

The first base model of SEM is a Multi-layer Perceptron (MLP). MLP is a type of artificial neural network that consists of layers of artificial neurons, and the input is processed layer by layer [14]. The calculation of the output for each neuron is as follows [14]:

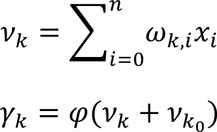

where *v_k_* is the net output of neuron *k*, with *x*_1_, *x*_2_, . . ., *x_n_* being the input from the last layer, and *ω*_*k*,1_, *ω*_*k*,2_, . . ., *ω_k,n_* being the weights on the input. *γ_k_* is the output of the neuron, where *v*_*k*_0__ represents the bias, and φ(.) refers to the activation function. And the MLP part is trained with Rooted Mean Squared Error (RMSE) loss:

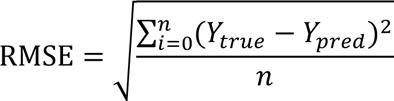

where *n* is the size of the input dataset, *Y_true_* is the true value of copy number, *Y_pred_* is the value of the predicted copy number.

In addition to MLP, the K-mer counts data would undergo a dimension reduction by the Principal Component Analysis (PCA) algorithm, implemented by the *PCA* function in scikit-learn with *n_components* set to 100, and then be input to the several other machine learning models, including Support Vector Machine (SVM) [46], Random Forest [47], K-nearest Neighbors [48], and Gradient Boosting [49]. Linear Regression, Ridge Regression [50], and Lasso Regression [51] were tested as meta-models which accept the output of MLP, PCA and machine learning models and make the final prediction. All machine learning algorithms were implemented by the scikit-learn module (version 0.23.2). After the initial testing, SVM was selected among the base-models and Ridge Regression among the meta-models (Figure 2c).

**Figure 2.**
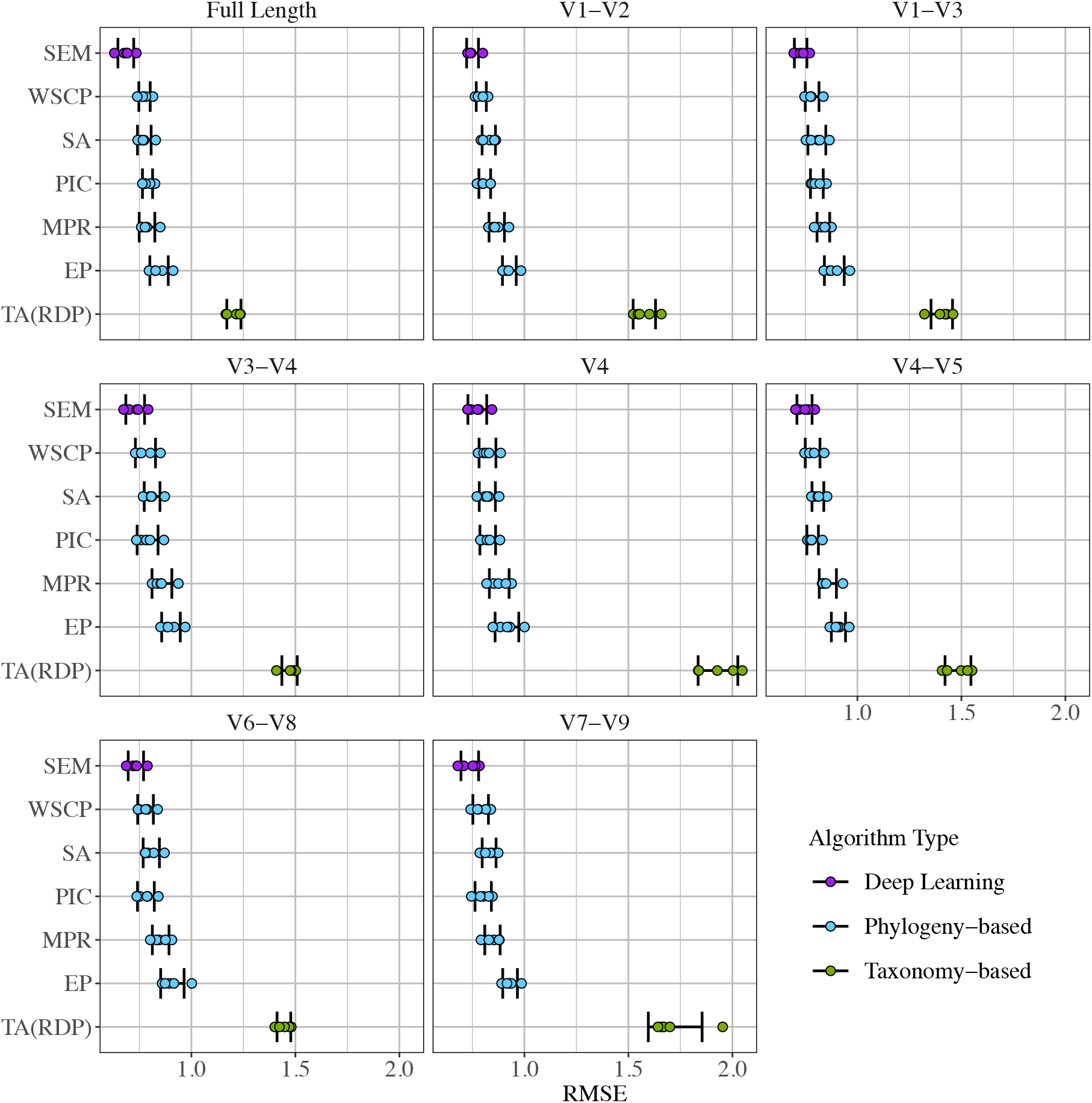
Algorithm performance comparisons at 16S full-length and commonly sequenced subregions. EP: empirical probabilities; MPR: Sankoff’s maximum-parsimony; PIC: phylogenetic independent contrast; SA: Subtree Averaging; WSCP: weighted-squared-change parsimony; TA(RDP): taxonomic aggregation based on RDP database; SEM: stacked ensemble model.

#### 2.3.3. Parameter Optimization

The parameter of SEM was optimized through the *SMAC4BB* function in the SMAC3 package (version 1.4.0) [52] for 500 iterations. For the MLP component, hyperparameters including the number of dense layers, hidden layer size, activation functions, and learning rate were tuned. The number of dense layers of the model was tested in the range of [1, 7], hidden layer size from the range of [1, 1024], activation functions are selected among Rectified Linear Unit (RELU), Exponential Linear Unit (ELU), Gaussian Error Linear Unit (GELU), Scaled Exponential Linear Unit (SELU), and Linear, and the learning rate was tested in the range of [0.00005, 0.1]. The mathematic formulae of the activation functions are demonstrated in the Supplementary Methods.

The final MLP architecture (Figure 2d) contains a dense block of 6 hidden layers, with neuron numbers of 489, 926, 645, 929, 582, and 82 respectively, and the activation functions are GELU, ReLU, ReLU, ELU, GELU, and Linear respectively. For SVM, the *kernel* value was chosen from “rbf”, “sigmoid”, and “linear”, *gamma* was tested between “scale” and “auto”, and *C* was tested as a float in the range of [0.3, 0.9] or as an integer in the range of [1, 100]. For Ridge, the value of *alpha* was tested as a float in the range of [0.3, 0.9] or as an integer in the range of [1, 100]. After tuning, the optimized parameters are *kernel = ‘rbf ’*, *gamma = ‘auto’*, *C = 11*, and *alpha = 49* (a complete list of parameters is shown in Table S2).

The resulted final model is named the Artificial Neural Network Approximator for 16S rRNA Gene Copy Number (ANNA16).

#### 2.3.4. Other Deep Learning Models Tested

In addition to SEM, this study also developed and evaluated a Transformer model and a Residual Multi-layer Perceptron model for 16S GCN prediction. Details about these two models are described in the Supplementary Methods and Figure S1 & S2. However, neither model achieved perform comparable to the SEM and they were not selected.

### 2.4. Mock Community Test

To assess model performance, an *in silico* test study was designed based on two commercially available mock communities [53], one even community (ATCC, MSA-2003) and one staggered community (ATCC, MSA-1001) (Figure 5, Table S3). Both mock communities consist of ten common bacterial strains, but of different ratios. This study retrieved genome sequences of the ten strains from NCBI, and copy number data from rrnDB (version 5.7). An arbitrary total cell number (1 × 10^5^ cells) were assigned to each community and the cell number for each strain was calculated based on the known/targeted fraction.

To evaluate the accuracy of community profile prediction by different tools, Bray-Curtis Dissimilarity [54] was calculated between the true cell fraction and the estimated cell fraction using the *vegdist* function in the vegan (version 2.5-7) package.

### 2.5. Model Explanation

In this study, SHAP a unified measure of feature importance that quantifies individual feature contributions to a model prediction, was utilized to explain DNA loci’s contributions [26]. Based on collational game theory [55] and the classical Shapley method [56], SHAP views each feature as a player. The final contribution of a feature is the sum of all the marginal contributions across various subsets. Positive and negative SHAP values indicate positive and negative contribution to the model’s prediction, respectively. In this study, each K-mer is considered a feature, and its SHAP value is calculated using the *KernelExplainer* function in shap module (version 0.41.0) in Python, treating each single prediction as an independent game.

This study randomly sampled 10% of the whole dataset for SHAP value analysis (N = 2758). As a result, a list of 2,758 SHAP values was assigned to every K-mer to represent the K-mer’s overall contribution. These SHAP values were mapped back to the 16S rRNA gene sequence according to the position of the corresponding K-mers to investigate if there is a positional pattern in contribution. What is more, to study the relationship between contribution and nucleotide conservation in a position, the rate of nucleic substitution, insertion, and deletion of each position was calculated using aligned 16S rRNA gene sequences, with *E. coli* (accession number: GCF_002953035.1) as the reference, to represent how conserved a position. The upper boundary of the plot of SHAP values versus the insertion rate was fitted with the elu function to explore how ANNA16 weighs one insertion. To represent the upper boundary, the insertion rate was first clustered with the *KMeans* function in the scikit-learn package into 50 ranks. For each rank, the mean insertion rate and max SHAP values were calculated. Adam optimizer in the tensorflow package were used to fit the elu function with a learning rate of 0.01 and 2000 iterations.

## 3 RESULTS

This study evaluated the performance of SEM in comparison with other algorithms for full-length 16S GCN correction. Figure 2 and Table 1 summarize the results of 5-fold cross-validation. Among all algorithms, taxonomic aggregation methods (TA) perform the worst. TA based on RDP or Greengenes reference database exhibit a mean RMSE in copies/genome 1.18 (SD = 0.0356) and 2.52 (SD = 0.0388), respectively. Due to the much higher mean RMSE of TA with Greengenes, only results from TA with RDP are shown in Figure 2. All phylogeny-based algorithms perform better than taxonomic aggregation, with WSCP the best phylogeny-based algorithm. SEM, the deep learning model, outperforms all other algorithms with the lowest mean RMSE 0.685 copies/genome, with SD = 0.0379. Its error range on the 16S full-length is significantly lower than all other algorithms (p < 0.01).

**Table 1.** Performance of all algorithms at 16S full-length and subregions (Mean RMSE ± SD, *p-*value).

|  | Full Length | V1-V2 | V1-V3 | V3-V4 | V4 | V4-V5 | V6-V8 | V7-V9 |
| --- | --- | --- | --- | --- | --- | --- | --- | --- |
| SEM | 0.685 ± 0.0379 | 0.751 ± 0.0285 | 0.727 ± 0.0301 | 0.730 ± 0.0450 | 0.774 ± 0.0453 | 0.746 ± 0.0364 | 0.733 ± 0.0368 | 0.737 ± 0.0423 |
| EP | 0.845 ± 0.0443<br>$p < 0.001$ | 0.928 ± 0.0331<br>$p < 0.001$ | 0.889 ± 0.0476<br>$p < 0.001$ | 0.902 ± 0.0447<br>$p < 0.001$ | 0.916 ± 0.0569<br>$p < 0.01$ | 0.910 ± 0.0340<br>$p < 0.001$ | 0.909 ± 0.0563<br>$p < 0.001$ | 0.931 ± 0.0354<br>$p < 0.001$ |
| MPR | 0.787 ± 0.0375<br>$p < 0.01$ | 0.867 ± 0.0370<br>$p < 0.001$ | 0.836 ± 0.0305<br>$p < 0.001$ | 0.858 ± 0.0477<br>$p < 0.01$ | 0.879 ± 0.0475<br>$p < 0.01$ | 0.858 ± 0.0413<br>$p < 0.001$ | 0.852 ± 0.0400<br>$p < 0.001$ | 0.846 ± 0.0370<br>$p < 0.01$ |
| PIC | 0.789 ± 0.0243<br>$p < 0.001$ | 0.810 ± 0.0281<br>$p < 0.01$ | 0.805 ± 0.0307<br>$p < 0.01$ | 0.789 ± 0.0503<br>$p < 0.05$ | 0.824 ± 0.0377<br>$p < 0.05$ | 0.785 ± 0.0277<br>$p < 0.05$ | 0.781 ± 0.0404<br>$p < 0.05$ | 0.802 ± 0.0393<br>$p < 0.05$ |
| SA | 0.774 ± 0.0329<br>$p < 0.01$ | 0.829 ± 0.0321<br>$p < 0.01$ | 0.805 ± 0.0430<br>$p < 0.01$ | 0.811 ± 0.0379<br>$p < 0.01$ | 0.822 ± 0.0388<br>$p = 0.054$ | 0.810 ± 0.0283<br>$p < 0.01$ | 0.807 ± 0.0391<br>$p < 0.01$ | 0.830 ± 0.0334<br>$p < 0.01$ |
| WSCP | 0.775 ± 0.0273<br>$p < 0.05$ | 0.793 ± 0.0242<br>$p < 0.05$ | 0.782 ± 0.0330<br>$p < 0.05$ | 0.779 ± 0.0483<br>$p = 0.067$ | 0.823 ± 0.0405<br>$p = 0.054$ | 0.785 ± 0.0354<br>$p = 0.062$ | 0.779 ± 0.0377<br>$p < 0.05$ | 0.790 ± 0.0375<br>$p < 0.05$ |
| TA(RDP) | 1.18 ± 0.0356<br>$p < 0.001$ | 1.58 ± 0.0538<br>$p < 0.001$ | 1.41 ± 0.0517<br>$p < 0.001$ | 1.47 ± 0.0370<br>$p < 0.001$ | 1.93 ± 0.0954<br>$p < 0.001$ | 1.48 ± 0.0622<br>$p < 0.001$ | 1.44 ± 0.0329<br>$p < 0.001$ | 1.73 ± 0.129<br>$p < 0.001$ |
| TA(Greengenes) | 2.52 ± 0.0388<br>$p < 0.001$ | 2.08 ± 0.122<br>$p < 0.001$ | 2.63 ± 0.147<br>$p < 0.001$ | 1.85 ± 0.0939<br>$p < 0.001$ | 2.12 ± 0.0590<br>$p < 0.001$ | 2.00 ± 0.0961<br>$p < 0.001$ | 2.19 ± 0.232<br>$p < 0.001$ | 2.54 ± 0.434<br>$p < 0.001$ |

EP: empirical probabilities; MPR: Sankoff's maximum-parsimony; PIC: phylogenetic independent contrast; SA: Subtree Averaging; WSCP: weighted-squared-change parsimony; TA(RDP): taxonomic aggregation based on RDP database; TA(Greengenes): taxonomic aggregation based on Greengenes database; SEM: Stacked Ensemble Model.

Furthermore, algorithm performance was evaluated and compared for commonly amplified 16S subregions (Figure 2). Compared to the performance on full length, in general all algorithms produce larger RMSE on subregions. SEM displays relatively consistent performance on subregions, ranging from 0.731 (V1-V3) to 0.774 (V4) copies/genome. Notably, except for V3-V4, V4, and V4-V5 of WSCP and V4 of SA, the error range of SEM on subregions is significantly lower than all other algorithms in all subregions (p < 0.05). These findings indicate SEM outperforms EP, MPR, PIC, and TA with RDP or Greengenes, while partially outperforms SA, and WSCP.

The prediction results produced by SEM from the first cycle of cross-validation are presented in Figure 3 and Figure S3. The figure shows a strong linear relationship between the predicted values and the true values, with density plot showing similar data distribution patterns. However, it is worth noting that SEM tends to underestimate the GCN when the true value is greater than 14. This issue is further discussed in Section 4.

**Figure 3.**
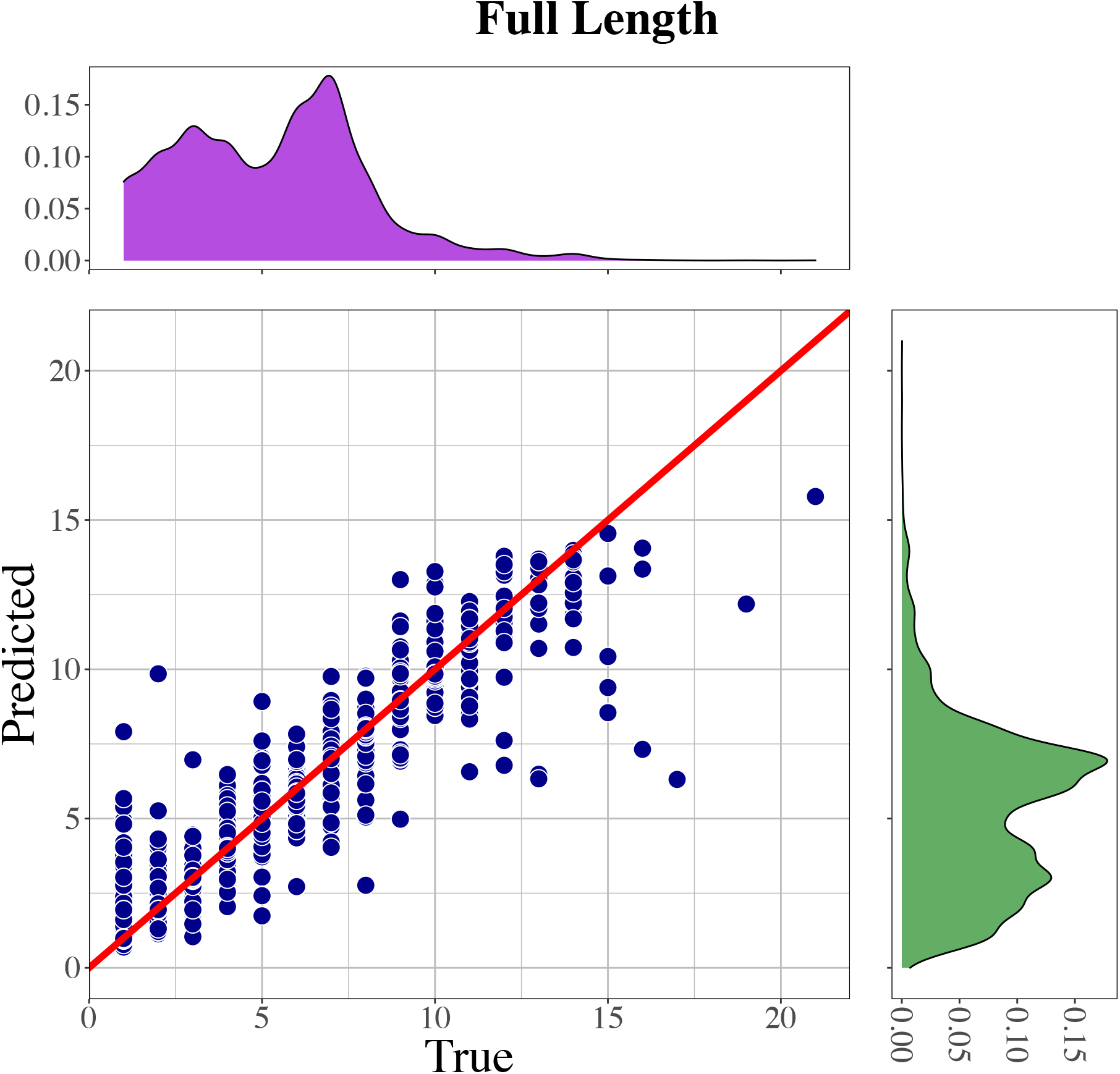
Prediction by SEM on 16S full-length in the first cycle of 5-fold cross-validation. The red line is a diagonal line illustrating to which extent the predicted values are derived from the true values. The density plot filled in violet shows the distribution of the true values, and the green density plot shows the distribution of the predicted values.

Due to SEM’s superior performance, a GCN prediction tool, ANNA16, is developed based on SEM and compared with other available tools in the final test, the newly published data in rrnDB version 5.8 (Figure 4 and Table 2). Because PICRRUSt2 has a much higher RMSE (i.e., larger than 3.4 copies/genome for all regions), it is excluded from Figure 4 and the following case study. ANNA16 achieves the lowest RMSE among all the tools examined, with the smallest RMSE (0.683) at the full-length and the highest RMSE (0.780) at V4. In contrast, the RMSE of CopyRighter ranges from 1.822 (V4-V5) to 2.181 (V7-V9) copies/genome, rrnDB from 1.079 (V4-V5) to 1.207 (V4) copies/genome, PAPRICA from 0.773 (V3-V4) to 1.321 (V4-V5), and PICRUSt2 from 3.459 (V4-V5) to 4.731 (V6-V8). ANNA16 outperformed all other tools, demonstrating superior performance at all subregions and full-length, which indicated its superior ability to predict GCN in newly discovered species.

**Figure 4.**
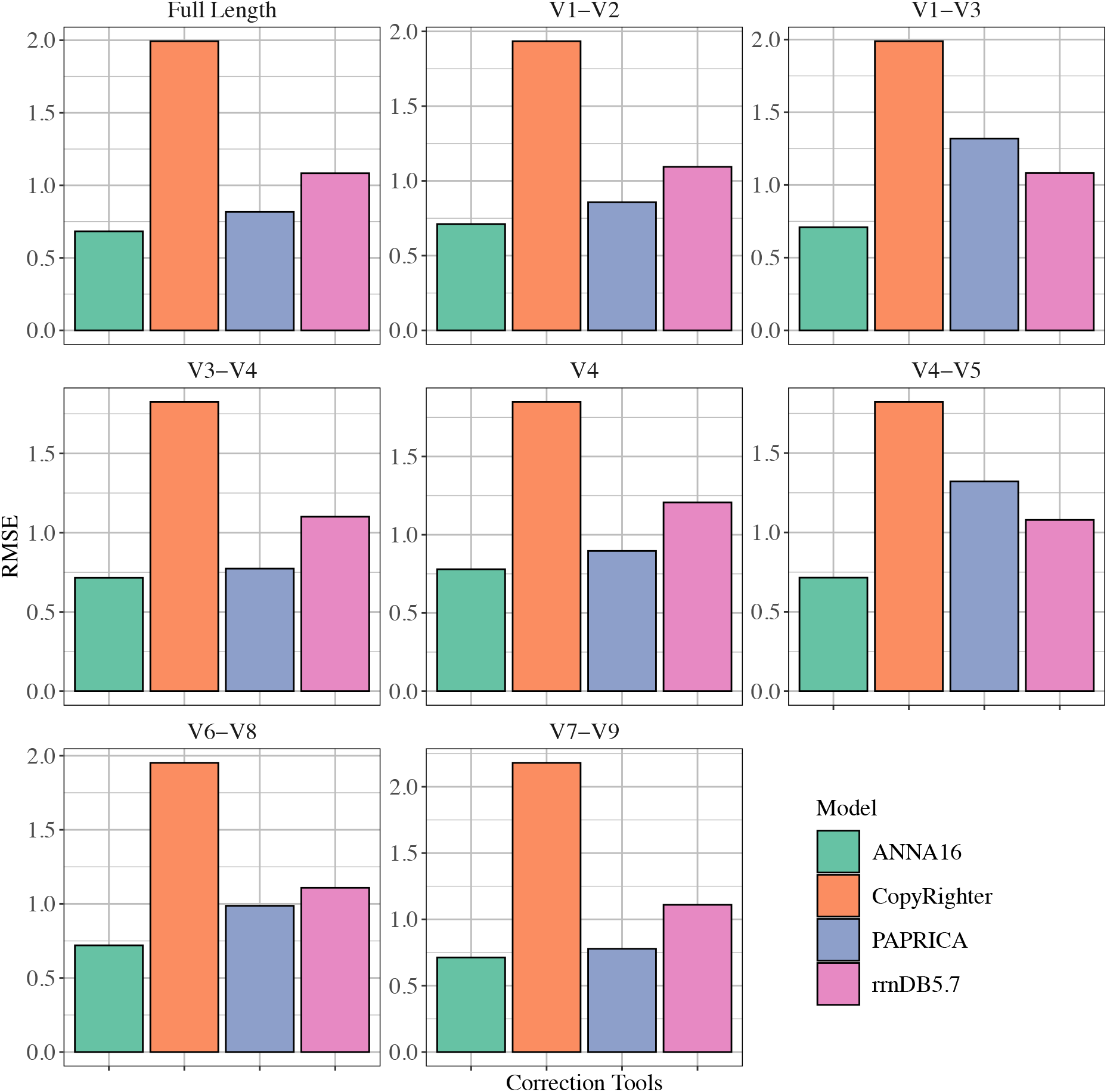
Comparison of performance on the final test dataset among CopyRighter, rrnDB5.7, ANNA16, and PAPRICA.

**Table 2.** Performance of 16S GCN Correction Tools in the Final Test (RMSE)

|  | Full Length | V1-V2 | V1-V3 | V3-V4 | V4 | V4-V5 | V6-V8 | V7-V9 |
| --- | --- | --- | --- | --- | --- | --- | --- | --- |
| CopyRighter | 1.994 | 1.934 | 1.989 | 1.825 | 1.848 | 1.822 | 1.952 | 2.181 |
| rrnDB5.7 | 1.083 | 1.094 | 1.082 | 1.101 | 1.207 | 1.079 | 1.109 | 1.110 |
| ANNA16 | 0.683 | 0.711 | 0.710 | 0.716 | 0.780 | 0.715 | 0.720 | 0.713 |
| PICRUSt2 | 4.198 | 4.353 | 4.175 | 4.393 | 3.807 | 3.459 | 4.731 | 4.677 |
| PAPRICA | 0.817 | 0.857 | 1.319 | 0.773 | 0.896 | 1.321 | 0.987 | 0.778 |

To demonstrate the performance of ANNA16 intuitively, a case study of *in silico* mock communities was conducted. Figure 5, Table S4, and Table S5 show uncorrected copy fraction, true cell fraction, and cell fraction (microbial profiles) estimated by ANNA16, rrnDB (version 5.7), CopyRighter, and PAPRICA. The difference in uncorrected copy fraction and true cell fraction demonstrates that GCN correction is necessary for obtaining an accurate microbiome profile. The Bray-Curtis Dissimilarity analysis shows that in general the dissimilarity values for the even community are considerably higher than those for the staggered community. The cell fraction estimated by ANNA16 has the smallest dissimilarity value for both even and staggered communities. For the even community, PAPRICA and CopyRighter exhibit very close dissimilarity values and are the second most accurate tools. For the staggered community, PAPRICA is the second most accurate tool, followed by rrnDB, but CopyRighter is the least accurate one. Overall, ANNA16 outperforms all the other tools in terms of similarity patterns with the true cell fraction.

**Figure 5.**
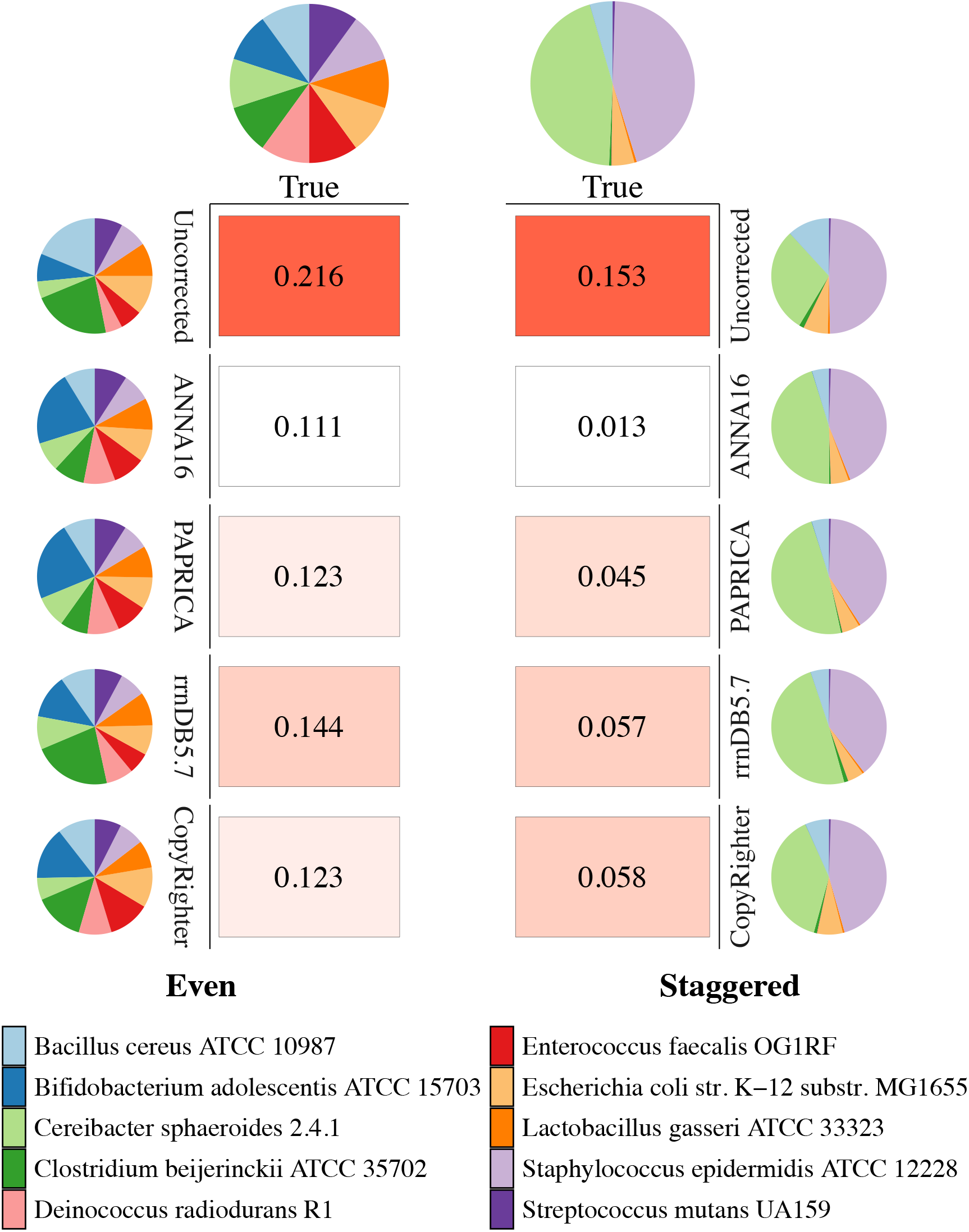
*In silico* test with the mock communities. Left panels: The composition of the even community estimated by uncorrected 16S gene copy fraction, ANNA16, PAPRICA, rrnDB (version 5.7), CopyRighter and their Bray-Curtis Dissimilarity with the cell fraction. The Dissimilarity values are presented as the heat map. Right panel: The composition of the staggered community estimated by the methods and their Bray-Curtis Dissimilarity with the cell fraction. Uncorrected: Uncorrected copy fraction; True: True cell fraction; ANNA16: Cell fraction estimated by ANNA16; rrnDB5.7: Cell fraction estimated by rrnDB (version 5.7); CopyRighter: Cell fraction estimated by CopyRighter; PAPRICA: Cell fraction estimated by PAPRICA. Even Community: All bacterial strains account for the same proportion. Staggered Community: The ratio of each bacterial strains is uneven.

To illustrate how ANNA16 makes its predictions, the shap module in Python is used to calculate the SHAP value of each K-mer, which represents the contribution of each K-mer to the prediction. Figure S4 displays the top 20 K-mers ranked by mean absolute SHAP value, while Figure 6 illustrates mean absolute SHAP values of each nucleic position across the full length 16S rRNA gene sequences. The pattern does not fully coincide with the hypervariable regions, which implies the distribution of high SHAP values does not strictly align with the traditionally identified hypervariable regions. To confirm this implication, the relationship between SHAP value and the rate of mutations (substitution, insertion, and deletion) is plotted in Figure 7. Figure 7a and 7b shows that no non-linear pattern could be observed between SHAP values and substitution or deletion, and the Pearson correlation coefficients indicate the linear association is also weak (cor=0.144 or cor=0.248). However, intriguingly, the insertion rate exhibits a non-linear relationship with SHAP values, with high SHAP values reaching its peak when the insertion rate is extremely low. These results imply ANNA16 mainly relies on rare insertions to predict GCN, instead of substitutions.

**Figure 6.**
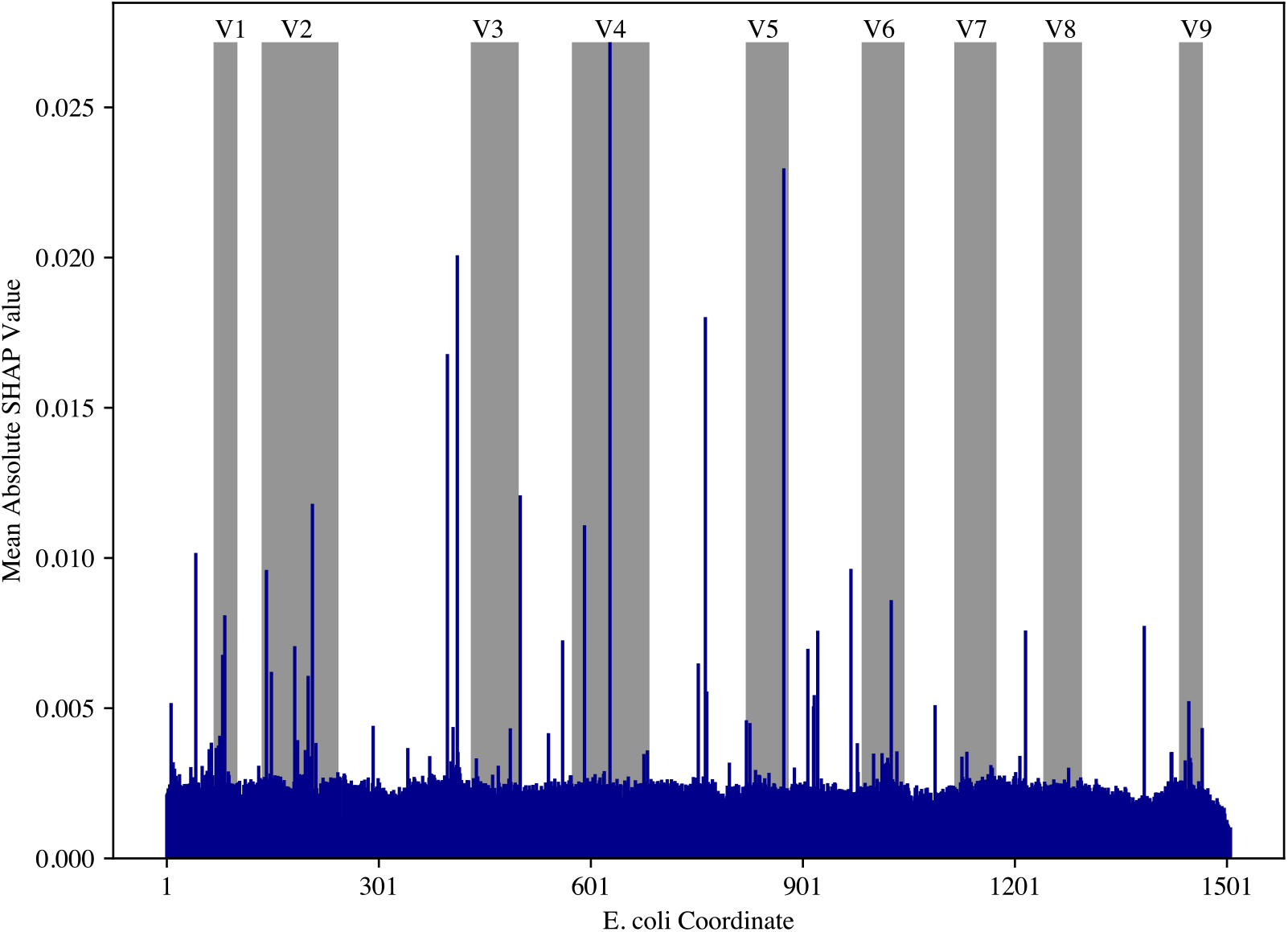
Mean absolute SHAP values across 16S rRNA gene full-length sequences. The mean absolute SHAP values were calculated using the aligned sequences, and then converted to *E. coli* coordinates with *E. coli* (accession number: GCF_002953035.1) as the reference.

**Figure 7.**
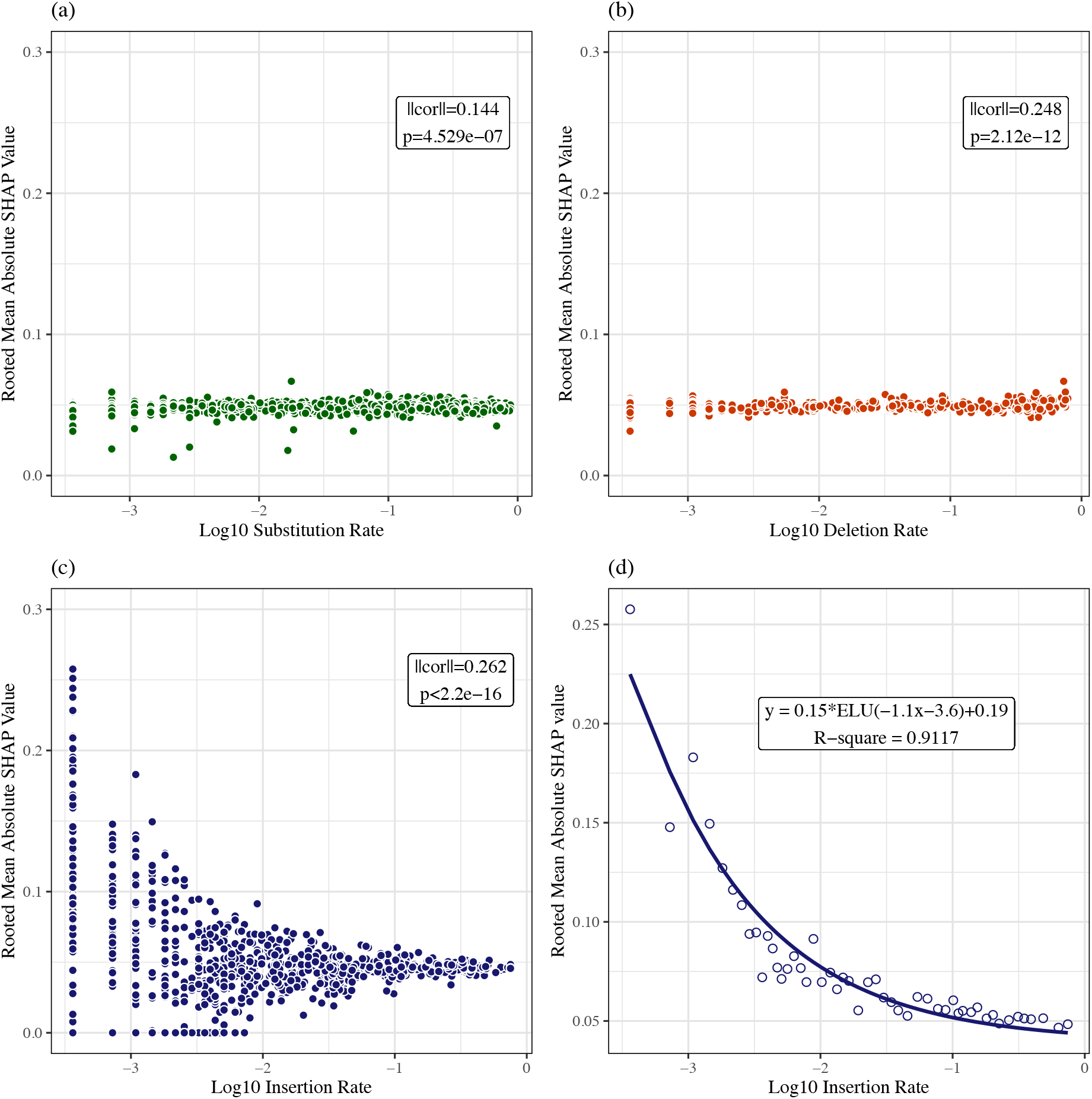
Relationship of SHAP values and the rate of mutations on each position of the aligned 16S rRNA gene sequences. (a) Log10 Substitution Rate vs. Rooted Mean Absolute SHAP Value. (b) Log10 Deletion Rate vs. Rooted Mean Absolute SHAP Value. (c) Log10 Insertion Rate vs. Rooted Mean Absolute SHAP Value. (d) The upper boundary of Log10 Insertion Rate vs. Rooted Mean Absolute SHAP Value, fitted by the elu function.

## 4 DISCUSSION

This study introduces ANNA16, a novel method for predicting 16S GCN based on 16S DNA sequence using a deep learning model. This novel approach outperforms currently available taxonomy- and phylogeny-based algorithms in accuracy, excelling in predicting GCN not only for the 16S full-length sequences but also in all commonly amplified subregions.

### 4.1. Strengths of ANNA16

Compared with existing GCN correction tools, the ANNA16 method owns their strengths and meanwhile avoids the drawbacks. Existing GCN correction tools can be classified into two categories based on their inference strategies: pre-computation or real-time phylogenetic placement. On the one hand, pre-computational tools (i.e., rrnDB & CopyRighter) compute the GCN value for every taxon in the database beforehand, and the tools retrieve the GCN values from the database when a query is given. This strategy has the advantage of short inference time, but is also heavily relay on the reference database, which can lead to errors if used with databases having differences in nomenclature. For example, the SILVA database [57] merges *Allorhizobium*, *Neorhizobium*, *Pararhizobium*, and *Rhizobium* as one genus, named *Allorhizobium-Neorhizobium-Pararhizobium-Rhizobium*, but in RDP these are independent genera.

On the other, real-time phylogenetic placement tools (i.e., PAPRICA and PICRUSt2) place a query sequence into their built-in phylogenetic trees to predict the GCN value at runtime. This strategy leads to better prediction performance than pre-computational tools, as shown in Figure 2 and 4, but requires much longer inference time because phylogenetic placement can be computationally extensive. However, the performance of ANNA16 can even be better than real-time phylogenetic placement tools, as shown in Figure 2 and 4. Compared with these tools, ANNA16 exhibits even better performance than real-time phylogenetic placement tools, avoids database-dependence by predicting GCN values directly from 16S rRNA gene sequences, and maintains a relatively short inference time because the prediction can be accelerated by GPUs.

### 4.2. Explaining ANNA16

After proving the excellence of ANNA16, an intriguing question this study contemplates is how a deep learning model interprets the intricate biological/evolutionary information contained within the 16S DNA sequence. Interestingly, Figure 6 and Figure 7 suggest ANNA16’s approach is highly different from existing methods.

First, the peaks of high SHAP values across the 16S full-length sequence are not aligned with the nine hypervariable regions. As shown in Figure 6, although there are peaks of high SHAP values at the V1, V2, V4, and V5 regions, yet the conserved regions before V1 and between V2, V3, V4, V5, and V6 also contain some loci that are significantly contributory. Those conserved regions even seem more contributory than the remaining variable regions (i.e., V3 and V6-V9). These results indicate the concept of variability is not equivalent to informativeness for ANNA16.

Second, SHAP values shows no association with nucleic substitutions. Previous studies mainly use nucleic substitutions to infer evolutionary information. For example, the fraction of nucleic substitutions between two sequences are commonly used as a representation for evolutionary distance [58], or the substitution rate was used as the metric for studying the positional variability in 16S rRNA gene [59]. However, as shown in Figure 7a, the rooted SHAP values remain at a constant low level (about 0.05) despite the variation in the substitution rate. This means implies that substitutions may not be so important for ANNA16, as every substitution makes the similarly low degree of contribution regardless of the prevalence.

Given the specialty of ANNA16, SHAP values was plotted against the deletion and insertion rate (Figure 7b & 7c) to further explore how ANNA16 perceives the information in DNA sequences. The deletion rate demonstrates a similar pattern with SHAP values as the substitution rate, where the rooted SHAP values also remain near 0.05. However, an intriguing L-shaped trend emerges between the insertion rate and the rooted SHAP values. High SHAP values aggregate at the positions with the lowest end of the insertion rate, whereas in other areas, SHAP values also maintain the constant level of 0.05. This pattern suggests that insertions of extremely low frequency are the most informative for ANNA16. Each rare insertion may only exist in a limited number of taxonomic lineages, so they could serve as identifiers of the corresponding lineages for ANNA16. What is more interesting, the upper boundary of Figure 7c appears to be highly similar with the elu activation function used in ANNA16. After using the elu function to fit the upper boundary, 91.17% variability in the SHAP values could be explained (Figure 7d). In general, these results imply that ANNA16 might understand one DNA sequence by screening for rare insertions.

One potential explanation for these trends in Figure 6 and Figure 7 is that some neurons in the MLP part of ANNA16 are differentiated for detecting the rare insertions, and they are assigned with high weights by other neurons. Similar to biological neurons, artificial neurons also receive input signals and fire an output to other neurons if the stimulus reaches a certain threshold. Hypothetically, some differentiated neurons might only fire when ANNA16 encounters one insertion whose rarity reaches the threshold (roughly 10^-2^ per site according to Figure 7d). This might result in the elu-shape of the upper boundary in Figure 7c, which places high SHAP values at positions with rare insertions and reduces the SHAP values of the remaining positions to a relatively low level.

These implications may explain why ANNA16 or MLP outperform conventional algorithms. In the aspect of information extraction, traditional algorithms have two drawbacks. First, these algorithms typically weigh every position equally by converting the variance in the sequence-to-sequence similarity, which can underrepresent informative positions. Second, especially for phylogeny-based algorithms, some information could be wasted while inferring the phylogeny as most tree-building methods would ignore alignment gaps [58, 60-63]. Previous studies have proved that indels are likely to experience few homoplasy and may produce more congruent phylogenetic trees if combined with substitutions [58, 60-63]. However, ANNA16 adopts a more nuanced approach, which not only considers the substitutions but also pays more attention to insertions. This allows ANNA16 to more effectively harness the information embedded in the 16S sequences, confirming the necessity of including indels in phylogenetics. Future studies could also consider turning from classic theory-based methods to machine learning, a data-driven and alignment-free approach, for extracting the evolutionary information from the sequences. The effectiveness of ANNA16 also highlights the significant potential of deep learning in genetics. Not only can deep learning models like ANNA16 accurately predict traits based on sequencing data, but by analyzing the contribution of each DNA feature, researchers can also establish connections between specific genes or loci and the trait of interest. For example, by combining deep learning with the SHAP methodology, previous studies have evaluated the contribution of genes for predicting tissue types [64], or the pathway-level contribution for the phenotype of diffuse large B-cell lymphoma [65].

### 4.3. Limitations & Future Studies

Although ANNA16 has shown great performance in 16S GCN prediction, there are several limitations. One limitation is its tendency to underestimate the GCN of species with high GCN. As shown in Figure 3, ANNA16’s predictions become increasingly underestimated when the true GCN is greater than or equal to 14. This limitation could be attributed to the imbalanced distribution of GCN in the currently available dataset (Figure S5). Sequences with GCN ≥ 12 account for only a small proportion of the whole dataset, and sequences with GCN > 15 are extremely rare. The model could be improved with more new sequence data with high GCN species, as new sequencing techniques are rapidly generating more genomic data. Similar implications are also discussed by [4, 9]. Alternatively, data imbalance methods, such as class re-balancing, information augmentation, and module improvement [66] could be applied to ANNA16 and other future models to address the issue.

Furthermore, transfer learning [67, 68] could also be explored as a potential solution. In the framework of transfer learning, a deep learning model is first pre-trained on one or more source tasks to capture general features or representation, and then is fine-tuned to apply the captured knowledge to target tasks [69]. To utilize this strategy, future studies could first collect a large amount of unlabeled 16S rRNA sequences and pre-train a model with an autoregression or autoencoder task, similar to Large Language Models like BERT [70] or GPT [71], and then fine-tune the model on the child task, 16S GCN prediction, which may yield more accurate results.

## 5 CONCLUSIONS

In conclusion, this study developed ANNA16 as a deep learning-based new bioinformatic tool for predicting the gene copy number of 16S rRNA. Compared to existing taxonomy-based and phylogeny-based algorithms, ANNA16 showed significantly lower error ranges at 16S full-length and all 16S subregions in the cross-validation. Furthermore, in the final test and mock community case study, ANNA16 outperformed existing tools such as rrnDB, CopyRighter, PICRUSt2, and PAPRICA. ANNA16’s simple structure also offers several advantages, including faster inference speed, independence of pre-computation, and less demand for computing resources. The SHAP analysis implies ANNA16 could utilize the information contained in the 16S rRNA gene sequences more effectively by not only considering substitutions, but also paying more attention to rare insertions. Future studies could focus on strategies such as pre-training the model on unlabeled data and addressing data imbalance to further decrease the error of copy number correction.

## LIST OF ABBREVIATIONS

ANNA16: Artificial Neural Network Approximator for 16S rRNA Gene Copy Number
ELU: Exponential Linear Unit
EP: Empirical Probabilities
GCN: Gene Copy Number
GELU: Gaussian Error Linear Unit
MLP: Multi-layer Perceptron
MPR: Sankoff’s Maximum Parsimony
PIC: Phylogenetic Independent Contrasts
RELU: Rectified Linear Unit
SA: Subtree Averaging
SELU: Scaled Exponential Linear Unit
SEM: Stacked Ensemble Model
SHAP: Shapley Additive exPlanation
TA: Taxonomic Aggregation
WSCP: Weighted Squared Change Parsimony

## DECLARATIONS

### Ethics approval and consent to participate

Not applicable.

### Consent for publication

Not applicable.

### Availability of data and materials

16S rRNA sequences and the GCN data analyzed during the current study are available in the rrnDB repository, https://rrndb.umms.med.umich.edu/. The codes and model files of ANNA16 are available at https://github.com/Merlinaphist/ANNA16. ANNA16 is deployed on Google Colab, https://colab.research.google.com/drive/1XwpTMCHSfTmzpHyKrmiD8aC8C_1nndUV#scrollTo=M0jLLsHC9W4M.

### Authors’ contributions

J.M. conceived the study. J.M. and T.C. analyzed the data and implemented the models. Y.L. and M.M. provided mentorship for the study. All authors contributed to the manuscript writing.

## Acknowledgments

The authors would like to thank Dr. Jiang Long for his kind comments on algorithm design.

