## Supplementary Materials for "Deep Learning for Predicting 16S rRNA Gene Copy Number"

### 1 SUPPLEMENTARY METHODS

#### 1.1.Mathematic Formulae of Activation Functions

$$RELU(x) = \begin{cases} 0, x \leq 0 \\ x, x > 0 \end{cases}$$

$$ELU(x, \alpha = 1) = \begin{cases} \alpha(e^x - 1), x \leq 0 \\ x, x > 0 \end{cases}$$

$$SELU(x, s = 1.05070098, \alpha = 1.67326324) = \begin{cases} s\alpha(e^x - 1), x \leq 0 \\ sx, x > 0 \end{cases}$$

$$GELU(x) = xP(X \leq x), where\ P(X) \sim N(0,1)$$

#### 1.2.Other Deep Learning Models Tested

In addition to the SEM model, this study also tested a Transformer-based model and a Residual Multi-layer Perceptron (ResMLP) model. The framework and architecture of these models is shown in Figure S1.

##### 1.2.1. Transformer

For the transformer-based model in Figure S1, we leveraged the original transformer encoder architecture with cosine learning rate decay [1], followed by global average pooling, and two dense layers. As shown in Figure S1c, the transformer encoder takes embedding and position encoding as input. For biological sequences, a dense embedding is considered more effective than sparse tokenization. Each sequence is thus represented as a matrix $\mathbf{M} \in \mathbb{R}^{n \times d}$, by embedding each 6-mer into a numerical vector of dimension *d* via the embedding layer. The cosine positional encoding method is used to incorporate both absolute and relative positional information by constructing the matrix **P** from **M** [2], where each position has:

$$p(2k+1, i) = \sin\,(i/1000^{2k/d})$$

$$p(2k+2, i) = \cos\,(i/1000^{2k/d})$$

The encoder includes multi-head self-attention, position-wise feed-forward network, residual connection, and layer normalization [2]. With the combination of these modules, the transformer encoder can process contextual information parallelly with the attention mechanism. For one single attention head, each token-embedding $e_i$ is taken as query $q_i$ to compute *scaled dot attention score* with all token-embedding in the same input sequence, i.e., $[e_1, e_2, \dots, e_L]$, as key *k* and value *v*. Formally, the output of one attention head is defined as follows:

$$\text{head}(\boldsymbol{q}_i, (\boldsymbol{k}_1, \boldsymbol{v}_1), \dots, (\boldsymbol{k}_L, \boldsymbol{v}_L)) \; = \sum_{j=1}^{L} \alpha(\boldsymbol{q}_i, \boldsymbol{k}_j)\boldsymbol{v}_j$$

$$\alpha(\boldsymbol{q}_i, \boldsymbol{k}_j) = \text{softmax}(\frac{{\boldsymbol{q}_i}^T \boldsymbol{k}_j}{\sqrt{d}})$$

Thus, for the whole input sequence S, the attention score from the multi-head mechanism is

$$\mathbf{M} = \text{Multihead(S)} = \text{Concat}(\text{head}_1, \text{head}_2, \dots, \text{head}_n)\mathbf{W}$$

where all **W** parameters are learnable for linear transformation. By learning different linear projections independently, it captures the combination of different representation subspaces. The intermediate representation is passed into the feed-forward network for further non-linear transformation with ReLU activation. The Layer Normalization is used here to avoid gradient explosion and accelerate convergence, and the Dropout method is to prevent over-fitting.

1.2.2. Residual Multi-layer Perceptron

ResMLP are MLP models utilizing Residual Connection architectures. In this architecture, skip connections are built to jump over two to three layers. Such design may help avoid the problem of vanishing gradients or degradation [3]. The ResMLP model is published in 2021 for image classification. It attains good accuracy without the presence of attention layer, bringing a surprise to the data science community [4].

The ResMLP model in this study is trained with Rooted Mean Squared Error (RMSE) loss:

$$RMSE = \sqrt{\frac{\sum_{i=0}^{n}(Y_{true} - Y_{pred})^2}{n}}$$

where $n$ is the size of the input dataset, $Y_{true}$ is the true value of copy number, $Y_{pred}$ is the value of the predicted copy number.

Hyper-parameter tuning is performed on the number of dense blocks, the number of dense layers in a dense block, dense layer size, and learning rate. Hidden layer size is selected from the list of (128, 256, 512, 1024). The learning rate is tested from 0.001 to 0.002, with 0.0002 being a step. The final architecture of ResMLP for full-length DNA sequence is shown in Figure S1d & S1e.

##### 1.2.3. Results and Discussion

The performance of the deep learning models is shown in Figure S2. Transformer exhibit a mean RMSE of 0.861 copies/genome (SD = 0.0643) and ResMLP is 0.726 copies/genome (SD = 0.0381). Interestingly, the performance of Transformer is significantly worse than ResMLP ($p < 0.01$) and SEM ($p < 0.001$). This result is unexpected because Transformer-based models, such as BERT [5] or GPT [6], have achieved great success in the field of Natural

Language Processing (NLP). One potential explanation is that, as the computing cost of the self-attention mechanism increases quadratically with sequence length, Transformer-based models are typically incapable of modeling extended sequences [7]. As a reference, in NLP, classical models normally take inputs below 512 or 1024 [5]. Given biological sequences are more complex than natural language and our input length exceeds one thousand, it's plausible that the Transformer-based model gives a worse performance. If future studies would like to utilize Transform-based models to study 16S rRNA gene sequences, apart from trimming the sequences, non-overlap K-mer encoding could also be considered. The sequence encoding method this study adopted for the Transform was first to overlappingly convert the sequences into K-mers (Figure 1a) and second convert the K-mers into the token vectors. This method would encode one 1,500 nt DNA sequence into a vector of 1,495 tokens. However, if the DNA sequences are non-overlappingly split into 5-mers, the 1,500 nt sequence would be converted as a vector of only 300 tokens. The latter method, as used by [8], could reduce the length of 16S rRNA gene sequences to a Transformer-tolerable length. Although this method may increase the difficulty for the model to learn the patterns in nucleic arrangements, it is still possible to achieve good performance given enough data [8].

### 2 SUPPLEMENTARY TABLES

**Table S1.** Primers for Extraction of the 16S full-length and subregions [9, 10].

| Regions | Forward Primers | | Reverse Primers | |
|---|---|---|---|---|
| Full Length | 27F | AGA GTT TGA TCC TGG CTC AG | 1492R | TAC GGY TAC CTT GTT ACG ACT |
| V1-V2 | 27F | AGA GTT TGA TCC TGG CTC AG | 338R | GCT GCC TCC CGT AGG AGT |
| V1-V3 | 27F | AGA GTT TGA TCC TGG CTC AG | 534R | ATT ACC GCG GCT GCT GG |
| V3-V4 | 341F | CCT ACG GGA GGC AGC AG | 785R | GAC TAC HVG GGT ATC TAA TCC |
| V4 | 515F | GTG CCA GCM GCC GCG GTA A | 806R | GGA CTA CHV GGG TWT CTA AT |
| V4-V5 | 515F | GTG CCA GCM GCC GCG GTA A | 926R | CCG YCA ATT YMT TTR AGT TT |
| V6-V8 | 939F | GAA TTG ACG GGG GCC CGC ACA AG | 1378R | CGG TGT GTA CAA GGC CCG GGA ACG |
| V7-V9 | 1115F | CAA CGA GCG CAA CCC T | 1492R | TAC GGY TAC CTT GTT ACG ACT |

**Table S2.** Parameters of subunits in SEM.

| **MLP Part** | | |
|---|---|---|
| **Architecture** | **Number of Neurons** | **Activation** |
| | 489 | GELU |
| | 926 | ReLU |
| | 645 | ReLU |
| | 929 | ELU |
| | 582 | GELU |
| | 82 | Linear |
| | 1 | Linear |
| **Learning Rate** | 0.000167703 | |
| **Batch Size** | 100 | |
| **Epoch** | 59 | |
| **Classic Machine Learning Part** | | |
| **Model** | **Parameters** | **Value** |
| PCA | n_components | 100 |
| SVM | kernel | rbf |
| | gamma | auto |
| | C | 11 |
| Ridge | alpha | 49 |

**Table S3.** Information of mock communities.

| Strain | GenBank ID | 16S copy number | Cell Counts (Even) | Cell Counts (Staggered) |
|---|---|---|---|---|
| *Bacillus cereus* ATCC 10987 | NC_003909.8 | 12 | 100,000 | 44,800 |
| *Bifidobacterium adolescentis* ATCC 15703 | NC_008618.1 | 5 | 100,000 | 400 |
| *Clostridium beijerinckii* ATCC 35702 | NC_009617.1 | 14 | 100,000 | 4,500 |
| *Deinococcus radiodurans* R1 | NC_001263.1 | 3 | 100,000 | 400 |
| *Enterococcus faecalis* OG1RF | NC_017316.1 | 4 | 100,000 | 400 |
| *Escherichia coli* str. K-12 substr. MG1655 | NC_000913.3 | 7 | 100,000 | 44,800 |
| *Lactobacillus gasseri* ATCC 33323 | NC_008530.1 | 6 | 100,000 | 4,500 |
| *Cereibacter sphaeroides* 2.4.1 | NZ_AKVW01000001.1 | 3 | 100,000 | 44,800 |
| *Staphylococcus epidermidis* ATCC 12228 | NC_004461.1 | 5 | 100,000 | 44,800 |
| *Streptococcus mutans* UA159 | NC_004350.2 | 5 | 100,000 | 4,500 |

**Table S4.** Uncorrected, true, and estimated abundance of strains in even community.

| Strain | Uncorrected | True | ANNA16 | CopyRighter | rrnDB |
|---|---|---|---|---|---|
| *Bacillus cereus* ATCC 10987 | 0.1875 | 0.1000 | 0.0840 | 0.1055 | 0.0972 |
| *Bifidobacterium adolescentis* ATCC 15703 | 0.0781 | 0.1000 | 0.2160 | 0.1473 | 0.1238 |
| *Clostridium beijerinckii* ATCC 35702 | 0.2188 | 0.1000 | 0.0883 | 0.1408 | 0.2202 |
| *Deinococcus radiodurans* R1 | 0.0469 | 0.1000 | 0.0923 | 0.0919 | 0.0763 |
| *Enterococcus faecalis* OG1RF | 0.0625 | 0.1000 | 0.0901 | 0.1171 | 0.0599 |
| *Escherichia coli* str. K-12 substr. MG1655 | 0.1094 | 0.1000 | 0.0883 | 0.1124 | 0.0837 |
| *Lactobacillus gasseri* ATCC 33323 | 0.0938 | 0.1000 | 0.0975 | 0.0776 | 0.0942 |
| *Cereibacter sphaeroides* 2.4.1 | 0.0469 | 0.1000 | 0.0766 | 0.0616 | 0.0926 |
| *Staphylococcus epidermidis* ATCC 12228 | 0.0781 | 0.1000 | 0.0760 | 0.0713 | 0.0739 |
| *Streptococcus mutans* UA159 | 0.0781 | 0.1000 | 0.0909 | 0.0745 | 0.0781 |

**Table S5.** Uncorrected, true, and estimated abundance of strains in staggered community.

| Strain | Uncorrected | TRUE | ANNA16 | CopyRighter | rrnDB |
|---|---|---|---|---|---|
| *Bacillus cereus* ATCC 10987 | 0.1181 | 0.0448 | 0.0486 | 0.0668 | 0.0515 |
| *Bifidobacterium adolescentis* ATCC 15703 | 0.0004 | 0.0004 | 0.0011 | 0.0008 | 0.0006 |
| *Clostridium beijerinckii* ATCC 35702 | 0.0138 | 0.0045 | 0.0051 | 0.0090 | 0.0117 |
| *Deinococcus radiodurans* R1 | 0.0003 | 0.0004 | 0.0005 | 0.0005 | 0.0004 |
| *Enterococcus faecalis* OG1RF | 0.0004 | 0.0004 | 0.0005 | 0.0007 | 0.0003 |
| *Escherichia coli* str. K-12 substr. MG1655 | 0.0689 | 0.0448 | 0.0511 | 0.0712 | 0.0443 |
| *Lactobacillus gasseri* ATCC 33323 | 0.0059 | 0.0045 | 0.0057 | 0.0049 | 0.0050 |
| *Cereibacter sphaeroides* 2.4.1 | 0.2952 | 0.4479 | 0.4429 | 0.3899 | 0.4906 |
| *Staphylococcus epidermidis* ATCC 12228 | 0.4920 | 0.4479 | 0.4393 | 0.4514 | 0.3915 |
| *Streptococcus mutans* UA159 | 0.0049 | 0.0045 | 0.0053 | 0.0047 | 0.0042 |

### 3 SUPPLEMENTARY FIGURES

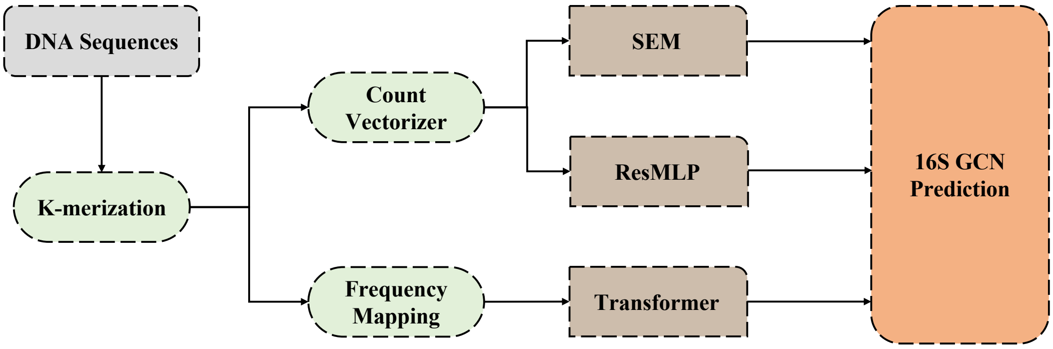


a) Framework of SEM, ResMLP, and Transformer

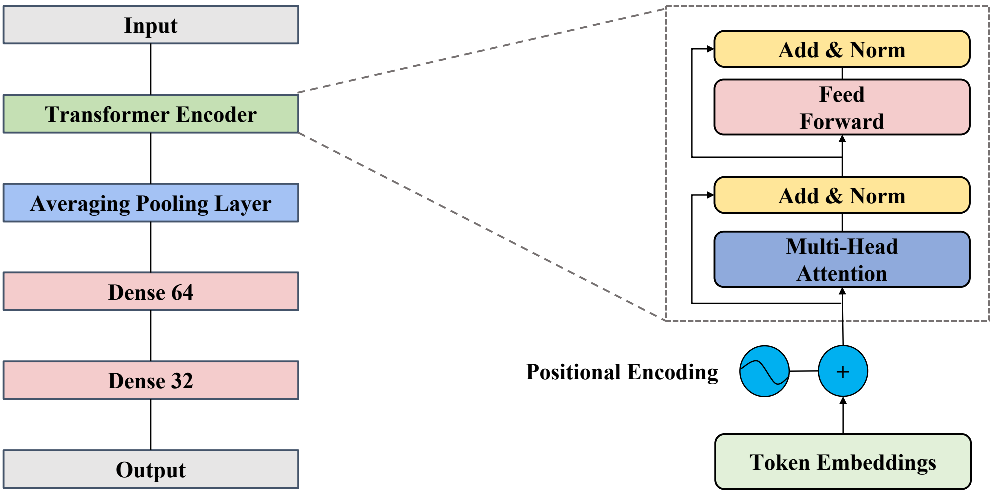


b) Architecture of the Transformer-based model

c) Architecture of the Transformer Encoder

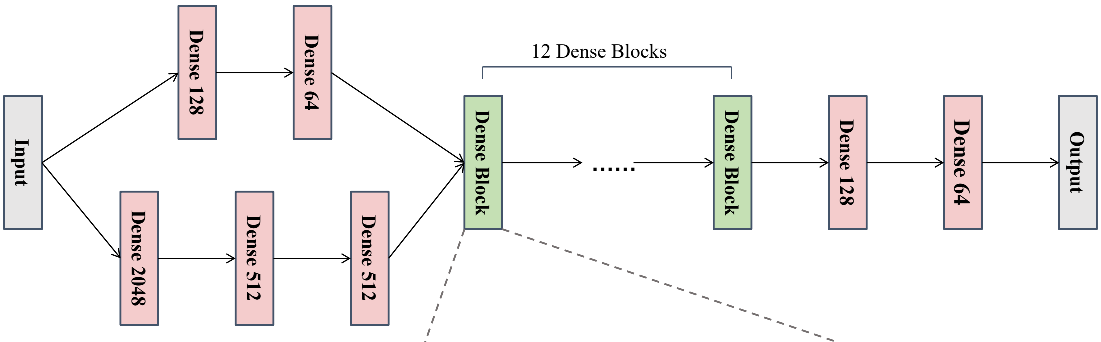


d) Architecture of the ResMLP model

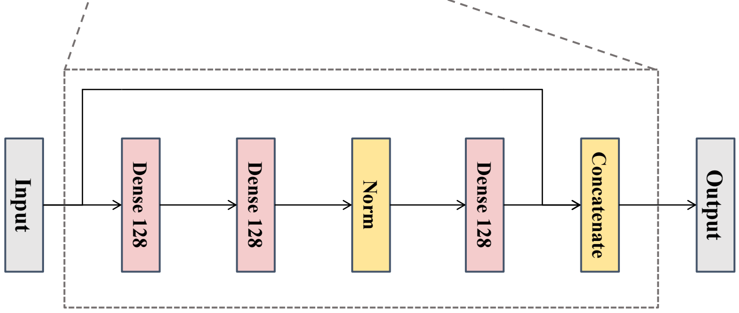


e) Architecture of a Dense Block

**Figure S1: Illustration of the Framework and Architecture of the Deep Learning Models.**

a) Framework of the SEM, ResMLP, and Transformer models. b) Architecture of the Transformer-based model. c) Architecture of the Transformer Encoder. d) Architecture of the ResMLP model. e) Architecture of a Dense Block inside the ResMLP model.

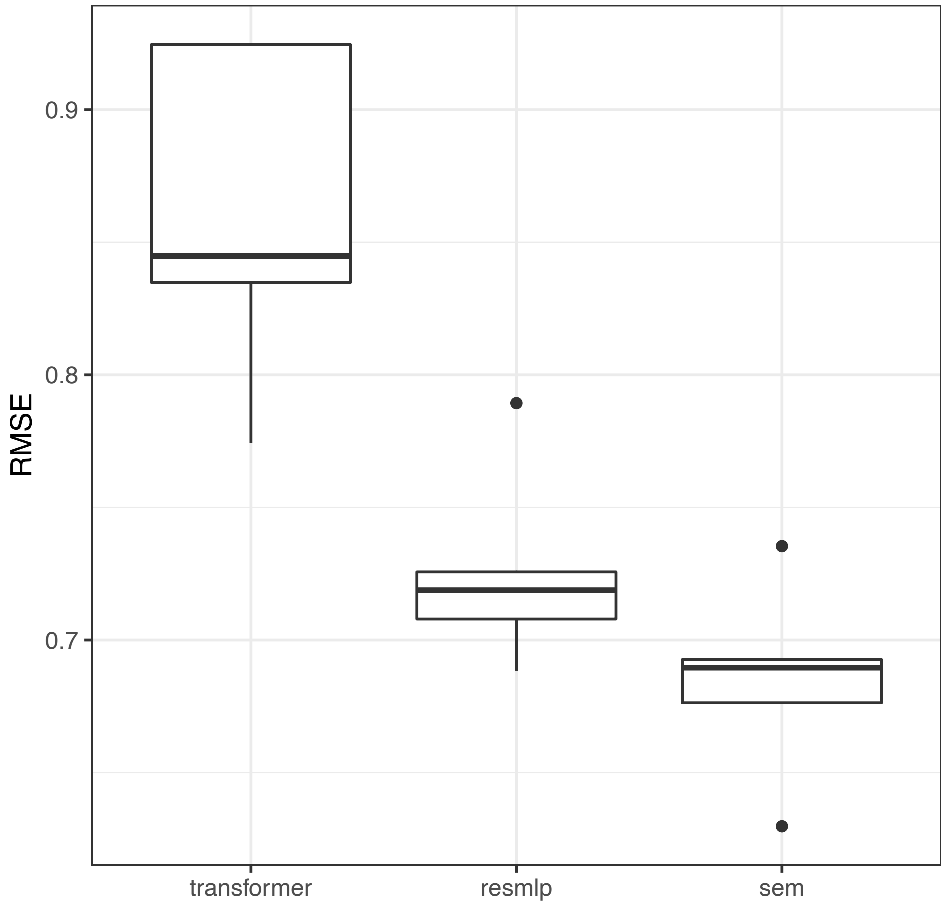


**Figure S2: Performance of the Deep Learning Models.**

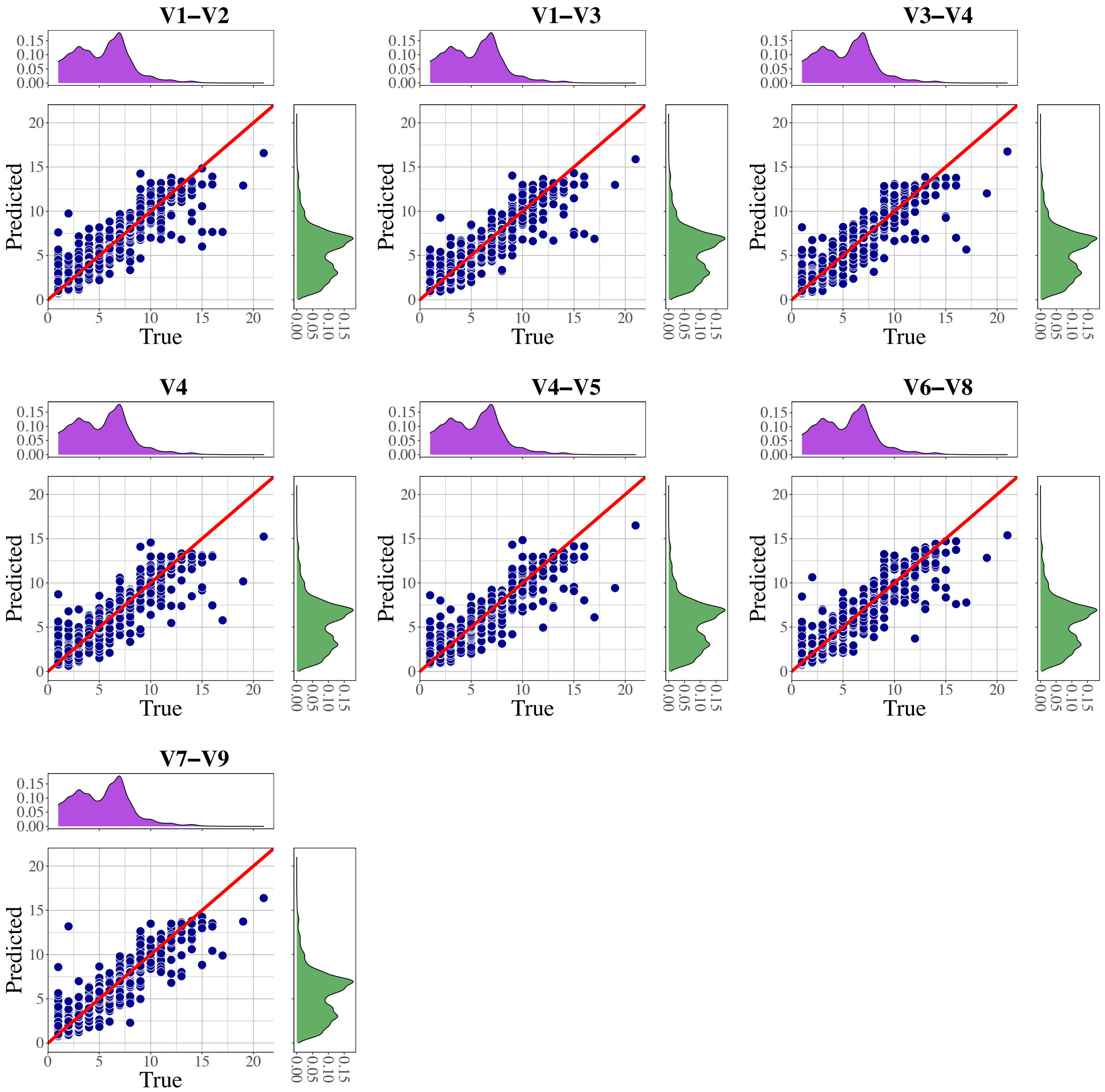


**Figure S3. Prediction by SEM on 16S Subregions in the First Cycle of 5-fold Cross-validation.** The red line is a diagonal line illustrating to which extent the predicted values are derived from the true values. The density plot filled in violet shows the distribution of the true values, and the green density plot shows the distribution of the predicted values.

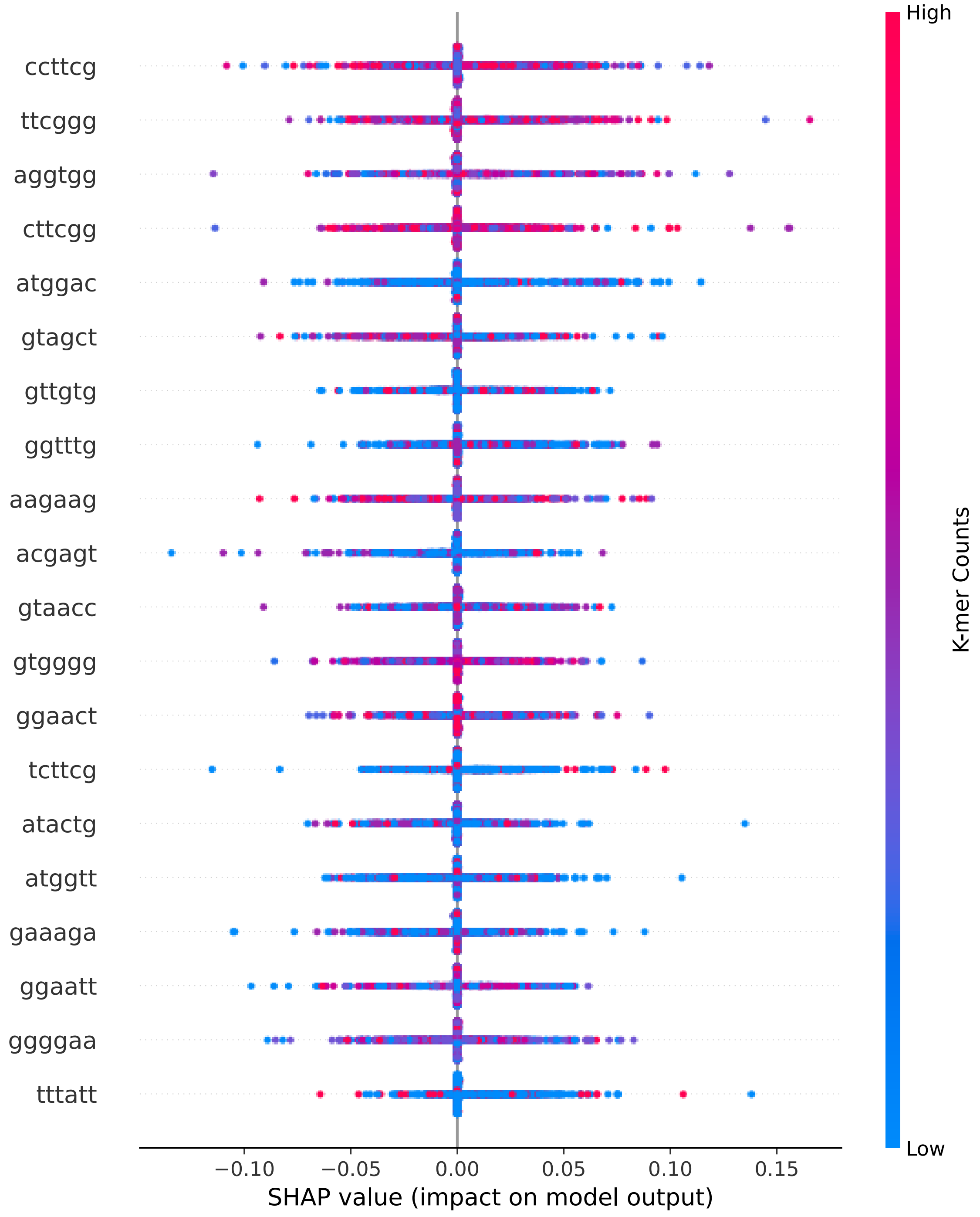


**Figure S4. SHAP values of top 20 K-mers ranked by mean absolute SHAP value.** Each point represents a SHAP value of a K-mer in a prediction.

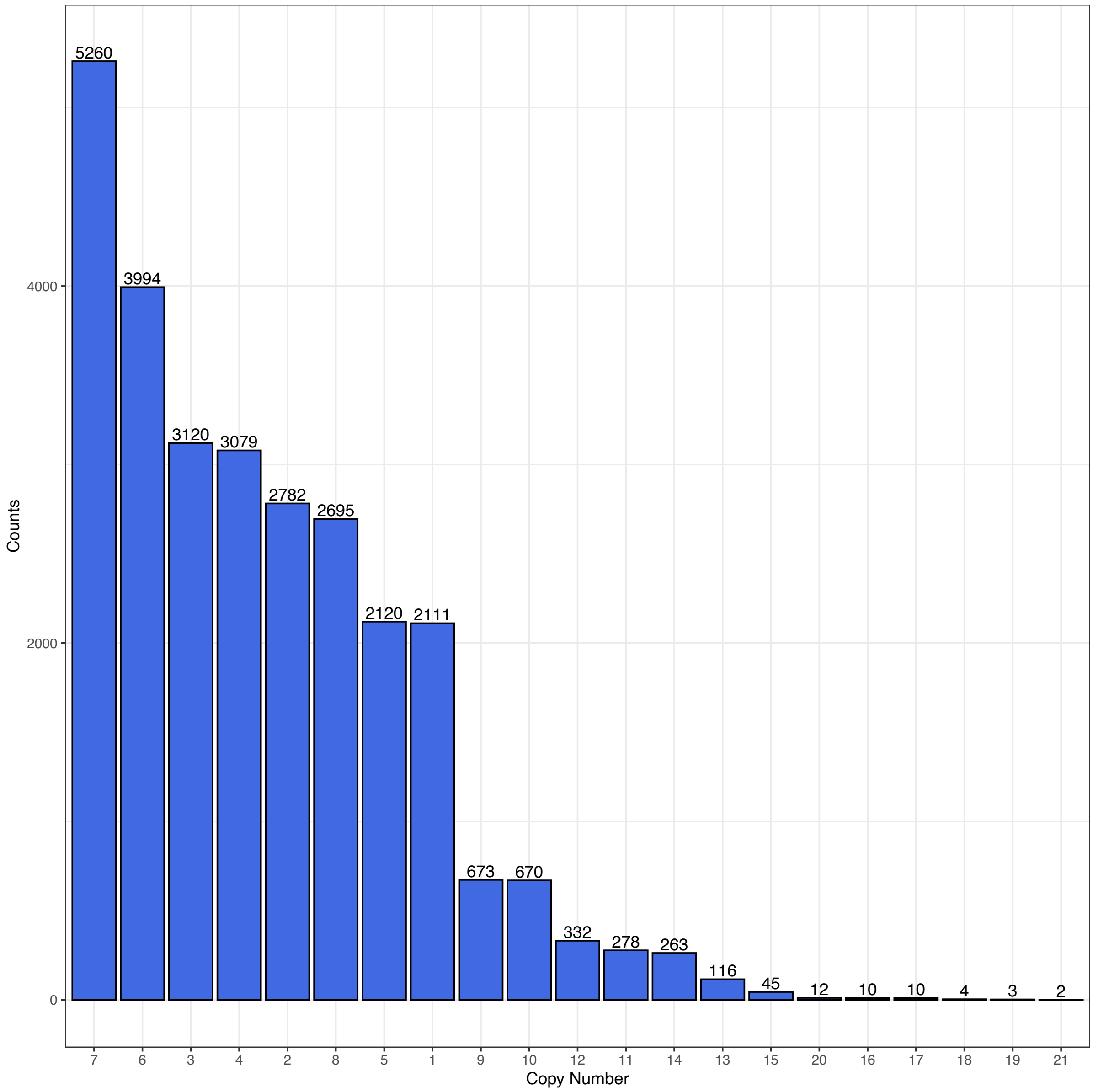


**Figure S5. Distribution of GCN in the whole dataset.**
